# Assessment of exposure to ionizing radiation in Chernobyl tree frogs (*Hyla orientalis*)

**DOI:** 10.1101/2021.05.17.443382

**Authors:** Pablo Burraco, Clément Car, Jean-Marc Bonzom, Germán Orizaola

## Abstract

Ionizing radiation can damage organic molecules, causing detrimental effects on human and wildlife health. The accident at the Chernobyl nuclear power plant (1986) represents the largest release of radioactive material to the environment. An accurate estimation of the current exposure to radiation in wildlife, often reduced to ambient dose rate assessments, is crucial to understand the long-term impact of radiation on living organisms. Here, we present an evaluation of the sources and variation of current exposure to radiation in breeding Eastern tree frogs (*Hyla orientalis*) males living in the Chernobyl Exclusion Zone. Total dose rates in *H. orientalis* were highly variable, although generally below widely used thresholds considered harmful for animal health. Internal exposure was the main source of absorbed dose rate (81% on average), with ^90^Sr being the main contributor (78% of total dose rate, on average). These results highlight the importance of assessing both internal and external exposure levels in order to perform a robust evaluation of the exposure to radiation in wildlife. Further studies incorporating life-history, ecological, and evolutionary traits are needed to fully evaluate the effects that these exposure levels can have in amphibians and other taxa inhabiting radio-contaminated environments.

## Introduction

Living organisms are constantly exposed to ionizing radiation. Cosmic rays, together with naturally occurring radioactive materials, generate low-level radiation known as background radiation (Sohrabi, 2013). Ionizing radiation has the capacity to damage organic molecules, including DNA, either directly by breaking DNA strains or through the generation of free radicals (Santivasi & Xia 2014). The main concern about the impact of ionizing radiation in wildlife is not generated by background radiation, but by the release of radioactive material to the environment due to human actions. These actions include nuclear weapons tests, mining of radioactive material, and accidents in nuclear facilities. The accidents in the nuclear power plants of Chernobyl (Ukraine, 1986) and Fukushima (Japan, 2011) represent the largest releases of ionizing radiation to the environment in human history. In order to reduce human exposure to radiation after these accidents, human settlement and normal activity were banned within certain areas, known as *Exclusion Zones*. In the absence of humans, wildlife becomes key study systems to examine the effects of the long-term exposure to ionizing radiation. Although the effects on health of the acute exposure to ionizing radiation were severe right after the Chernobyl accident (e.g. Geras’kin et al. 2008), there is still many uncertainties about the impact that chronic exposure to lower levels of ionizing radiation can have on wildlife (e.g. Møller and Mousseau, 2006, 2016; Beresford et al., 2020a,b). An accurate assessment of the exposure to ionizing radiation is needed to properly evaluate its consequences on the health of wild populations and across taxa.

The International Commission for Radiological Protection (ICRP) determined reference levels of radiation exposure called Derived Consideration Reference Levels (DCRLs), defined as a band of dose rate within which there is likely to be some chance of deleterious effects of ionizing radiation occurring to individuals (ICRP 2008). Different bands have been determined for a set of Reference Animals and Plants (RAPs; ICRP 2008). However, RAPs are limited to a few species, and DCRLs are defined mostly based on theoretical predictions or short-term laboratory procedures, thus they do not include the complexities of ecosystems, where organisms are often exposed to a wide array of fluctuating conditions and stressors (see e.g. Raines et al. 2020). Since RAPs are just single species, sometimes purely theoretical, that define entire animal or plant groups (e.g. “eusocial bee” defining all types of insects; ICRP 2008), they do not include either basic differences in species life styles, physiology, or morphology. Other thresholds levels have been proposed for organisms and ecosystems by different organizations (e.g. ERICA, FASSET, Environment Agency UK, Environment Canada; see summary in Garnier-Laplace et al. 2008), with standard levels above ICRP values, in most cases. More field studies, conducted in non-RAP organisms and under ecologically relevant scenarios, are clearly needed to understand the variability of exposure levels in wildlife.

More than three decades have passed since the Chernobyl nuclear power plant accident, a time that approximately corresponds to the half-life (i.e. the time required for a 50% reduction of the initial levels at the time of the accident) of ^90^Sr and ^137^Cs, the two main radioisotopes currently present in the Chernobyl Exclusion Zone (Beresford et al. 2010). Radiation levels in Chernobyl Exclusion Zone are now several orders of magnitude lower than at the time of the accident, and they are generated by a different array of radioisotopes (Beresford & Copplestone 2011). An accurate evaluation of current exposure to ionizing radiation in wildlife inhabiting the Chernobyl Exclusion Zone needs to consider the contribution of different radionuclides and radiation types (alpha, beta and gamma), and go beyond the use of portable dosimeters, which only estimate ambient dose rates, account only for gamma radiation, and do not distinguish between the contribution of different radioisotopes (Beresford et al. 2010). A detailed estimation of the current levels of exposure to ionizing radiation in wildlife living in radio-contaminated areas is crucial to assess the risk that radioactive substances can represent for these organisms, to provide a proper dosimetry context for understanding the effects (or lack of effects) of ionizing radiation in ecologically-realistic scenarios, and to estimate the accuracy of the proposed reference levels used in radiological assessment.

In this study, we examine the most important sources and the variation of the exposure to current ionizing radiation in breeding Eastern tree frog (*Hyla orientalis*) males living within the Chernobyl Exclusion Zone. ICRP uses a theoretical frog as a reference for predicting radiosensitivity in amphibians, for which a band of 40-400 μGy/h was defined within is likely to start detecting deleterious effects (ICRP 2008). The ERICA Tool, one of the most widely used software to assess radiological risk to terrestrial, freshwater and marine biota, used a default screening dose rate of 10 μGy/h for protecting organisms living in natural ecosystems (Brown et al. 2008). We use these references thresholds since they are widely used by the radioecology community, and are also two of the most conservative ones, with critical levels normally below other proposed references (Garnier-Laplace et al. 2008). Previous studies have reported a wide variation in the contribution of internal versus external exposure in amphibians, as well as differences in radioisotope contributions between species and areas (e.g. Beresford et al. 2020c). Here, we estimate dose rates in tree frogs collected during three consecutive breeding seasons (2016-2018) across the wide gradient of radioactive contamination currently present in the Chernobyl Exclusion Zone. In order to have a precise estimation of the current exposure to ionizing radiation in wild tree frogs, we not only quantified ambient dose rates, but also internal and external exposure to radiation in adult breeding frogs by integrating the activity of both ^90^Sr in bones and ^137^Cs in muscles. We expected to find a high contribution of internal dose rates and ^90^Sr (Beresford et al. 2020c), as well as high variability in dose rates across the Chernobyl Exclusion Zone. The understanding of the variability in radiation exposure in wild amphibians is critical for further evaluations of potential life-history and eco-evolutionary effects of radiation.

## Results

### Ambient dose rates across tree frog’s breeding habitats in the Chernobyl Exclusion Zone

Ambient dose rates measured at the twelve *H. orientalis* breeding localities sampled within the Chernobyl Exclusion Zone ranged from 0.07 to 32.40 μSv/h (Table 1). Six localities had ambient dose rates above 1 μSv/h and are located in areas commonly considered as highly contaminated (> 1000 kBq/m^2^ of ^137^Cs in 2018; Fig. 1). Six additional localities had ambient dose rates below 0.3 μSv/h (< 375 kBq/m^2^ of ^137^Cs in 2018; Fig. 1).

**Table 1.** Geographic coordinates (latitude and longitude), and current levels of environmental radiation (i.e. ambient dose rate) of the Eastern tree frog (*Hyla orientalis*) breeding localities included in the study. (* 7.61 μSv/h in 2017 and 2018; ** 1.09 μSv/h in 2017, 1.50 μSv/h in 2018, differences due to small changes in sampling areas within each locality and year).

| Locality | Code | GPS coordinates | Ambient dose rate ($\mu\text{Sv/h}$) |
| --- | --- | --- | --- |
| Vershina | VE | 51.4328, 30.0769 | 16.20 |
| Azbuchin | AZ | 51.4047, 30.1044 | 32.40 * |
| Muravka | MU | 51.4515, 30.0528 | 3.70 |
| Glyboke Hydro | GH | 51.4447, 30.0711 | 3.70 |
| Northern Trace | NT | 51.4567, 30.0486 | 2.51 |
| Dolzhikovo | DO | 51.4256, 30.1161 | 2.10 ** |
| Lubianka | LU | 51.3388, 29.7976 | 0.27 |
| Novosiolki | NO | 51.2195, 30.0430 | 0.13 |
| Zalesie | ZA | 51.2506, 30.1667 | 0.12 |
| Yampol | YA | 51.2119, 30.1899 | 0.10 |
| Glinka | GL | 51.2300, 29.9250 | 0.10 |
| Razjezheie | RA | 51.2786, 29.9050 | 0.07 |

**Figure 1.**
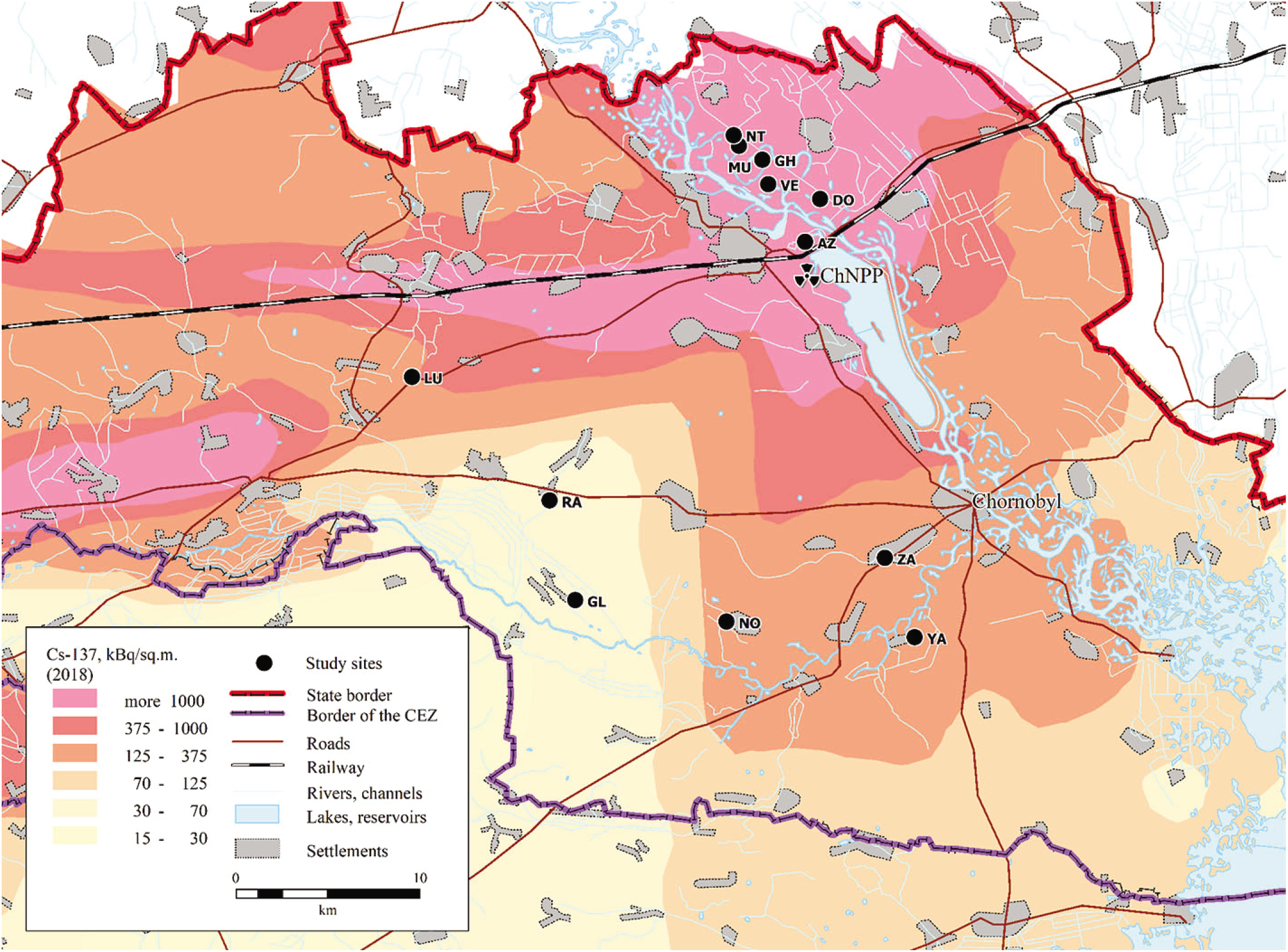
Map showing the localities where males of Eastern tree frog (*Hyla orientalis*) were sampled. The abbreviations refer to the locality name. Vershina (VE), Azbuchin (AZ), Muravka (MU), Glyboke Hydro (GH), Northern Trace (NT), Dolzhikovo (DO), Lubianka (LU), Novosiolki (NO), Zalesie (ZA), Yampol (YA), Glinka (GL), and Razjezzheie (RA; see Table 1 for details). The underlying ^137^Cs soil data (decay corrected to spring 2018) is derived from the Atlas of Radioactive Contamination of Ukraine (Intelligence Systems GEO, 2011).

### Radioactivity concentration in tree frog’s bones and muscles

Among the 226 male Eastern tree frogs (*Hyla orientalis*) examined, 65 individuals had activity concentrations below detection levels for ^90^Sr (29%), and 35 individuals for ^137^Cs (15%). In individuals where activity concentrations were above detection levels, ^90^Sr activity in bones ranged from 0.10 to 1156.98 Bq/g (fresh weight), which represents 0.0001 to 115.69 Bq/g of whole-body concentration (Supplementary data). Activity concentrations for ^137^Cs, measured in muscle tissue, ranged from 0.01 to 56.86 Bq/g (fresh weight), representing 0.0077 to 39.23 Bq/g of whole-body concentration (Supplementary data). The contribution of ^90^Sr to the total activity concentration of frogs living in localities with ambient dose rate > 1 μSv/h was, overall, two-fold higher than that of ^137^Cs (66% ^90^Sr contribution *versus* 33% of ^137^Cs contribution, on average).

### Dose rates of *H. orientalis* within Chernobyl Exclusion Zone

Total dose rates of *H. orientalis* males ranged between 0.01 and 39.35 μGy/h among individuals with activity rates above detection levels (Table 2, Fig. 2). Total dose rates varied substantially within and among localities, locality averages ranging from ca. 0 to 20 μGy/h (Table 2, Fig. 2). All sampled individuals had total dose rates below ICRP’s 40 μGy/h level for the reference frog (ICRP 2008), whereas ca. 20% (n = 46) had rates above ERICA’s 10 μGy/h screening dose rate limit for protecting ecosystems (Brown et al. 2008; Fig. 2). Internal dose rates ranged between 0.01 and 37.49 μGy/h, whereas external dose rates ranged between ca. 0 and 2.0 μGy/h (Table 2; Fig. S1). Internal and external dose rates were highly, and positively correlated (conditional R^2^ = 0.93; Fig. S1). For individuals living in areas where ambient dose rate was > 1 μSv/h (i.e. with individual dose rates above minimal detectable activities, see Methods), the contribution of internal dose rate to the total individual dose rate was always higher than the contribution of the external dose rate (83% of contribution of the internal dose rate, on average; χ^2^_(1,147)_ = 14.41, p < 0.001; Fig. 3). There was a highly significant and positive correlation between ambient dose rate and total individual dose rate (χ^2^(_1,226)_ = 15.21, p < 0.001, Estimate = 0.187; conditional R^2^ = 0.95; Fig. 4). The contribution of ^90^Sr represented, on average, 78% of the total dose rate of frogs living in localities with ambient dose rate > 1 μSv/h (Fig. 5), with a contribution of ^90^Sr to the internal dose rate six-fold higher than that of ^137^Cs (86% ^90^Sr contribution *versus* 14% of ^137^Cs contribution, on average, Fig. S2), despite only a 35% contribution of ^90^Sr to the external dose rate (Fig. S3, Supplementary data).

**Table 2.** Dose rates of breeding Eastern tree frog (*Hyla orientalis*) males captured within the Chernobyl Exclusion Zone (2016-2018). Data presented as mean value (range). Only localities sampled more than one year had variation in external dose rates. MDA: minimal detectable activity.

| Locality | Code | Sampled frogs (n) | Internal Dose Rate (μGy/h) | External Dose Rate (μGy/h) | Total Dose Rate (μGy/h) |
| --- | --- | --- | --- | --- | --- |
| Vershina | VE | 13 | 19.45 (34.77-7.65) | 1.51 | 20.96 (36.28-9.16) |
| Azbuchin | AZ | 46 | 13.31 (37.49-1.47) | 1.82 (2.00-1.69) | 15.13 (39.35-3-16) |
| Muravka | MU | 12 | 4.89 (7.72-2.92) | 0.58 | 4.81 (8.29-3.50) |
| Glyboke Hydro | GH | 10 | 4.72 (13.25-0.51) | 0.69 | 5.41 (13.94-1.19) |
| Northern Trace | NT | 18 | 2.46 (4.74-1.16) | 0.46 | 2.93 (5.20-1.63) |
| Dolzhikovo | DO | 49 | 2.25 (5.48-0) | 0.33 (0.34-0.33) | 2.58 (5.82-0.33) |
| Lubianka | LU | 5 | 0.09 (0.14-0.05) | 0.03 | 0.11 (0.17-0.10) |
| Novosiolki | NO | 12 | 0 (0.02-MDA) | MDA | 0.01 (0.02-MDA) |
| Zalesie | ZA | 12 | 0.02 (0.04-MDA) | 0.01 | 0.03 (0.05-0.02) |
| Yampol | YA | 14 | 0 (0.01-MDA) | 0.01 | 0.01 (0.02-0.01) |
| Glinka | GL | 25 | 0.01 (0.05-MDA) | MDA | 0.01 (0.05-MDA) |
| Razjezheie | RA | 10 | 0.11 (0.90-MDA) | MDA | 0.11 (0.91-MDA) |

**Figure 2.**
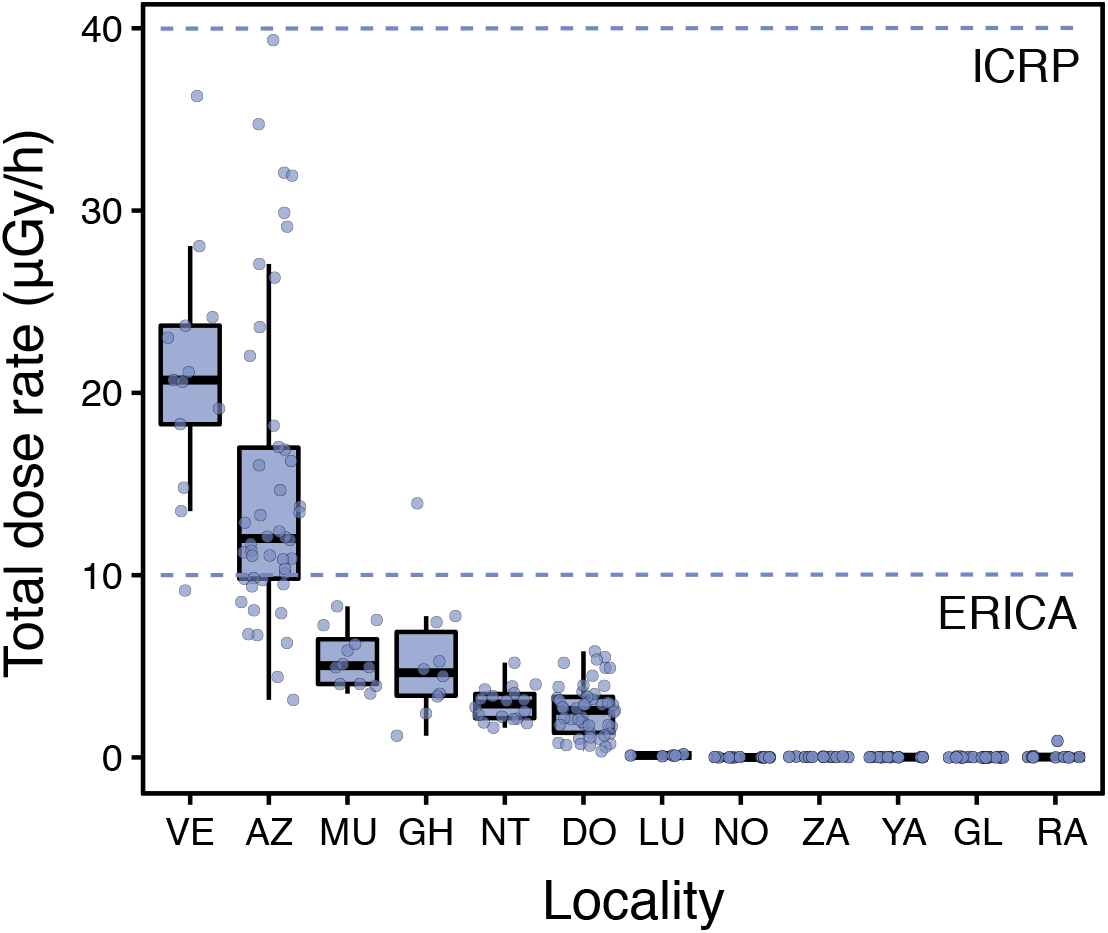
Total dose rates (μGy/h) of male breeding Eastern tree frogs (*Hyla orientalis*) living within Chernobyl Exclusion Zone. ICRP’s 40 μGy/h level for detecting damage on the reference frog, and ERICA’s 10 μGy/h screening level for protecting organisms within ecosystems are depicted with dotted lines. See Fig.1 for correspondence of locality.

**Figure 3.**
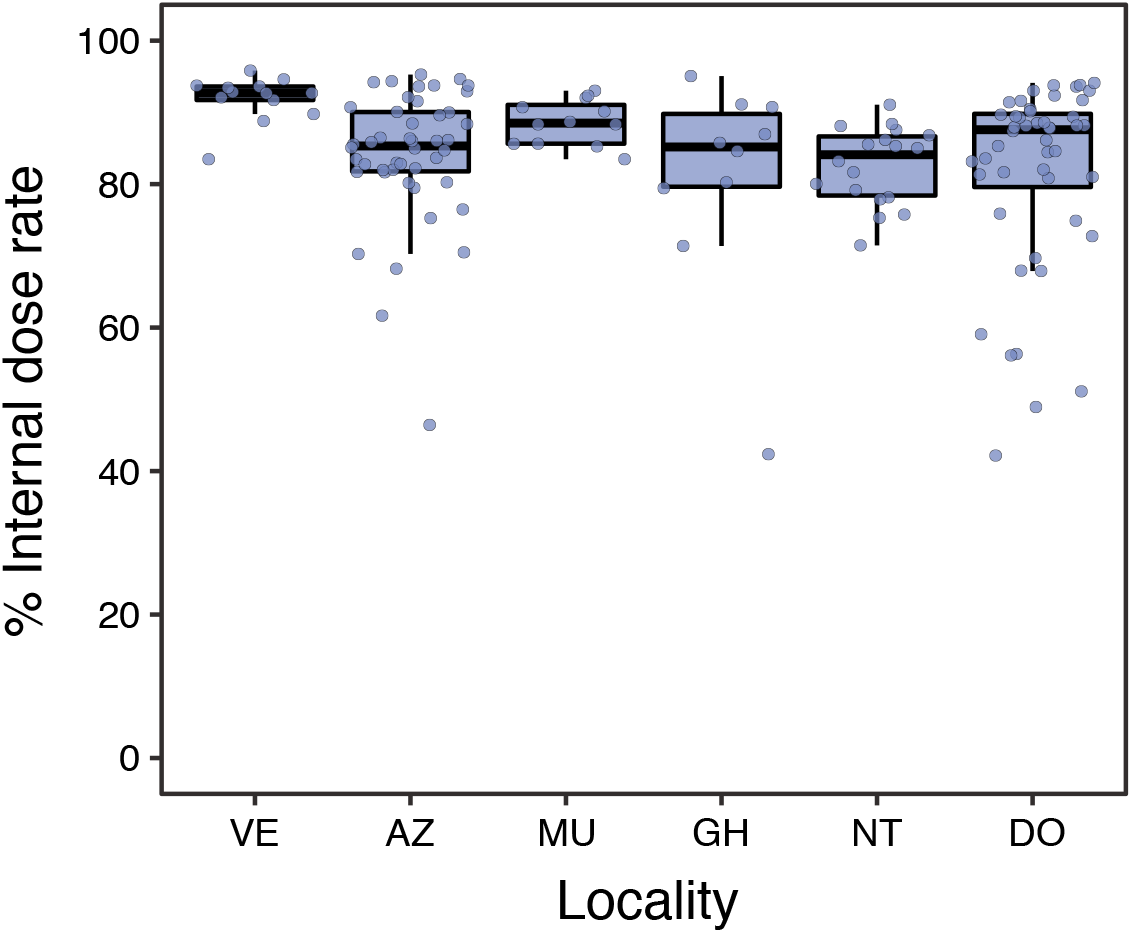
Contribution of internal dose rates (in percentage) to total individual dose rates of breeding Eastern tree frog (*Hyla orientalis*) males collected within the Chernobyl Exclusion Zone (only in individuals from localities with ambient dose rate > 1 μSv/h). See Fig.1 for correspondence of locality.

**Figure 4.**
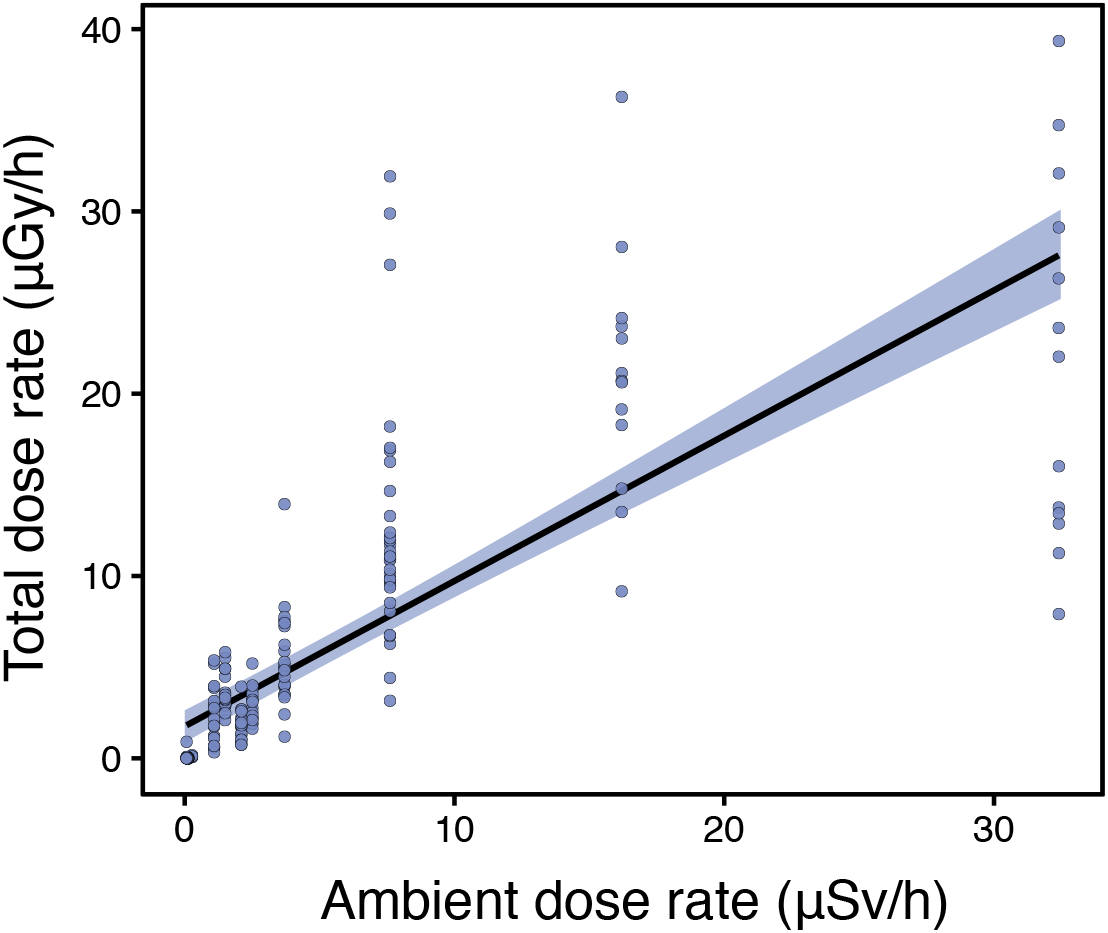
Correlation between ambient dose rates (in μSv/h) and total dose rates (in μGy/h) in breeding Eastern tree frog (*Hyla orientalis*) males living in the Chernobyl Exclusion Zone.

**Figure 5.**
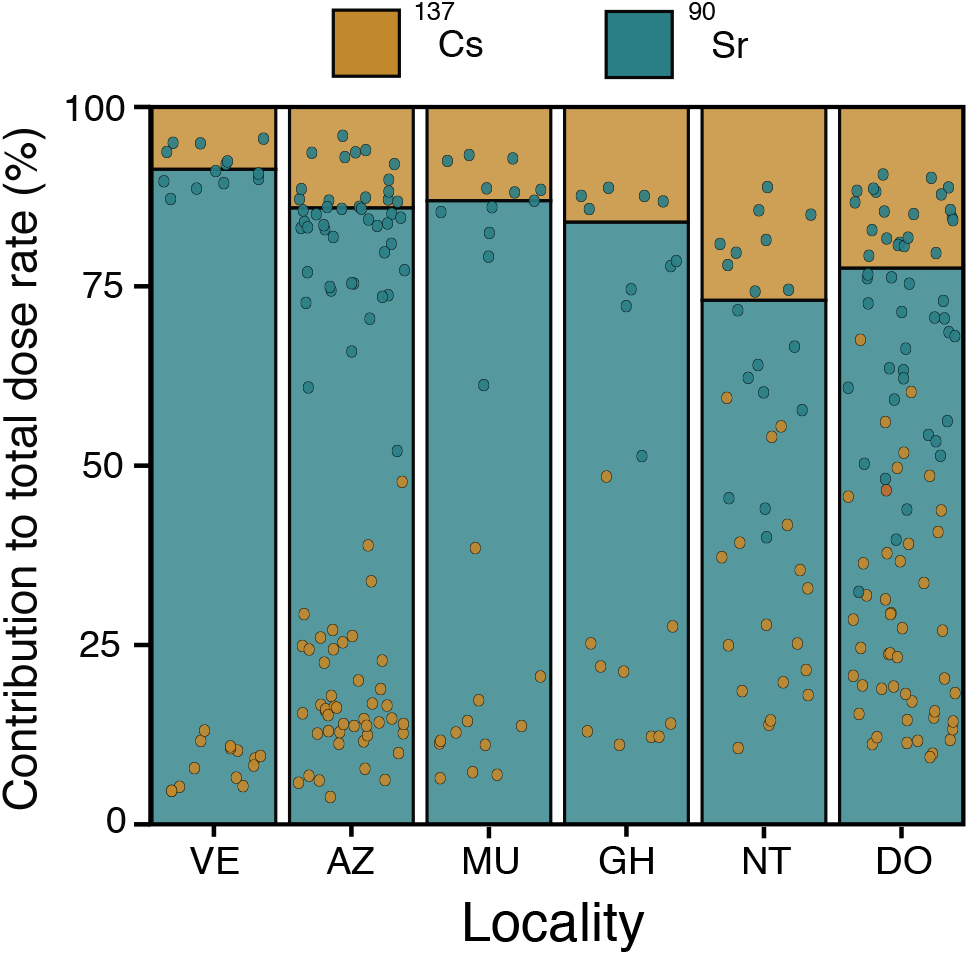
Contribution of ^90^Sr and ^137^CS (in percentage), to total dose rate in breeding Eastern tree frog (*Hyla orientalis*) males living in the Chernobyl Exclusion Zone (in individuals from localities with ambient dose rate > 1 μSv/h). Bars represent the locality average contribution for both isotopes, and points the contributions of each individual/isotope combination. See Fig.1 for correspondence of locality.

## Discussion

Our study shows that radiation exposure, in breeding males of the Eastern tree frog (*Hyla orientalis*) inhabiting across a wide gradient of radioactive contamination in the Chernobyl Exclusion Zone, is highly variable and, overall, below international thresholds for detecting damage. Individual dose rates varied substantially both at the inter- and intra-locality level. Total dose rates in Chernobyl tree frogs during the breeding season are dominated by internal, rather than external radiation levels, and are primarily a consequence of the doses of ^90^Sr in bones. Finally, although total individual dose rates were positively correlated with ambient dose rates, our data indicate that using only ambient dose rates will result in a poor estimation of the exposure to radiation in our study species as this parameter does not reflect the inter-individual variation in absorbed radiation.

Despite our study comprehensibly sampled frogs within twelve different localities and across the gradient of radioactive contamination in Chernobyl Exclusion Zone (Fig. 1), all the individuals presented total dose rates below the ICRP threshold level of 40 μGy/h (and also below other standards set by multiple organizations, see ICRP 2008, Garnier-Laplace et al. 2008). When using the more conservative 10 μGy/h screening level suggested by ERICA for protecting ecosystems (Brown et al. 2008), about 20% of the sampled *H. orientalis* were above this level, corresponding mainly to frogs collected in the two most radio-contaminated localities (AZ and VE, Fig. 2). Overall, these results suggest that three decades after the nuclear accident, exposure to radiation within the Chernobyl Exclusion Zone has dropped, in most cases, down below levels supposed to be damaging for these frogs during the breeding season (see Garnier-Laplace et al. 2008). Therefore, as a general prediction, negative effects of radiation are unlikely to be detected, except perhaps in the most radio-contaminated localities within Chernobyl (e.g. AZ and VE in our study, but see below). Anyway, these values need to be interpreted regarding the ecological characteristics that underlie our sampling design, as dose rates were measured on breeding individuals that expend a large amount of time in the interface between water/shoreline. A higher exposure is likely expected for frogs buried in the ground or leaf litter during the hibernation period, and therefore the within-year and lifetime variation in radiation levels deserves further exploration.

In our study, we examined ^90^Sr and ^137^Cs levels as sources of radiation for estimating internal and external dose rates. At present, ^90^Sr and ^137^Cs are the most abundant radioisotopes in the Chernobyl Exclusion Zone, whereas several radioisotopes with short half-life have already disappeared (e.g. ^131^I, ^132^Te, ^140^Ba; Beresford et al., 2010). However, other less abundant radionuclides such as ^241^Am, ^238^Pu, and ^239^Pu, are still present in the area and they might contribute to a fraction of the total dose rate accumulated by an organism (Beresford et al., 2020c). Nonetheless, previous studies conducted in the Chernobyl Exclusion Zone have reported a minimal contribution of these low-abundant isotopes to total dose rates in amphibians (Beresford et al., 2020c). Therefore, although our approach can slightly underestimate total dose rates in breeding tree frogs, we can consider that these differences should be minimal, and our dose rate estimates accurate.

Thresholds that determine radiation levels likely to cause damage are set without tests in ecological settings and can be inaccurate (see discussion in e.g. Raines et al. 2020). In our study system, further studies will determine whether current radiation levels experienced by Chernobyl tree frogs can negatively impact their life-history and eco-evolutionary dynamics (see e.g. Burraco et al. 2021). On this respect, there are important aspects that deserve further research. For example, we need to understand if adults of *H. orientalis* can have a higher sensitivity to radiation than the one predicted for the reference frog used by ICRP (ICRP 2008), as a consequence of differences in shape, size or life history between tree frogs, and the parameters considered when defining the ICRP theoretical reference frog. As commented above, dose rate can also vary across seasons and during the life-time of an individual. Furthermore, other ecological stressors such as diseases, parasites, or droughts, combined with small but still relevant effects of ionizing radiation can contribute to generate imbalances in the physiology and life-history of tree frogs at radiation levels defined as safe (Beresford et al. 2020a). Finally, effects of radiation currently observed can be a consequence of the impact of historical exposure to radiation, i.e. exposure to much higher radiation levels immediately after the accident (Beresford et al. 2020a). Therefore, differences across the radiation gradient within the Chernobyl area, but also between contaminated and non-contaminated localities outside the Exclusion Zone, may be linked to transgenerational carry-over effects induced by radiation in the past (transferred either by genetic and/or epigenetic mechanisms, Beresford et al. 2020a). Evaluating the relevance of these and other possible scenarios will improve our understanding on the effects that past and current exposure to ionizing radiation can have on wildlife health.

Our results also reveal that radioecology studies using only ambient radiation levels will inaccurately estimate the exposure of organisms to radiation (see e.g. comments in Beresford & Copplestone 2011, Beresford et al. 2012). We found a positive correlation between ambient dose rate and total individual dose rates in *H. orientalis*, suggesting that ambient radiation can be used to broadly define contamination areas for the species in Chernobyl. However, the high variation in individual total dose rates observed within each locality (i.e. for which ambient radiation is considered as a single value) indicates that using only ambient dose rates may lead to non-accurate estimates of the exposure to radiation experienced by each individual. Furthermore, previous studies on amphibians have revealed a large inter-specific variation in the contributions of internal and external dose rates. For example, internal dose rate represented ca. 40% of the total dose rate in moor frogs (*Rana arvalis*), ca. 50% in fire-bellied toads (*Bombina bombina*), and more than 70% in spadefoot toads (*Pelobates fuscus*), collected in the red forest area of Chernobyl Exclusion Zone (Beresford et al. 2020c). In other areas, internal dose rate was reported to have a minimal contribution to the total dose rate of moor frogs (*Rana arvalis*), collected in ponds of central Sweden within areas contaminated from the Chernobyl fallout (Stark et al. 2004). For other animal taxa, the contribution of internal dose rates can be as low as ca. 10% in bumblebees or ca. 20% in voles (*Microtus* spp.; Beresford et al. 2020c). Our study reports some of the largest contributions of internal dose rates reported for wildlife (83% internal contribution to total dose rate, Beresford et al. 2020c), and agrees with previous results in a similar species, the Japanese tree frog (*Hyla japonica*), examined in Fukushima and with internal dose rates contributing between 92-69% to the total dose rate (Giraudeau et al. 2018). Levels of ^90^Sr accumulated in the bones of *H. orientalis* contributed to most of the total individual dose rate (78%, on average), mostly due to its contribution to internal dose rate (86% on average, for frogs living in localities with ambient dose rate > 1 μSv/h). This also agrees with previous studies in Chernobyl reporting that ^90^Sr contributed between ca. 90% of the total dose rate in common toads (*Bufo bufo*) and spadefoot toads (*Pelobates fuscus*), and to a bit less than 60% in fire-bellied toads (*Bombina bombina;* Beresford et al. 2020c). Overall, ^90^Sr is the main source of total dose rates among Chernobyl wildlife (Beresford et al 2020c). Our results confirm the need to conduct detailed evaluations of internal exposure (i.e. internal dose rates) in order to precisely determine exposure levels in wildlife (Beaugelin-Seiller et al. 2003, Beresford et al 2020a,c).

Overall, this study presents a detailed evaluation of the variability of current exposure to radiation in breeding Eastern tree frogs (*H. orientalis*) living within the Chernobyl Exclusion Zone. Our study reveals the need to estimate total individual dose rates (i.e. including both internal and external exposure), and to evaluate the most common radioisotopes in order to accurately assess wildlife exposure to radiation. Dose rates, in our study species, are below widely used ICRP bands and most other proposed thresholds (ICRP 2008; Garnier-Laplace et al. 2008), whereas only 20% of the quantified dose rates were above ERICA screening levels for protecting ecosystems (Brown et al. 2008). However, many uncertainties remain around the estimation of these thresholds (e.g. Garnier-Laplace et al. 2008, Raines et al 2020), and therefore detailed studies incorporating life-history and eco-evolutionary variability are needed in order to properly evaluate the status of this species and other wildlife inhabiting Chernobyl.

## Methods

### Field sampling and laboratory procedures with Hyla orientalis

We used the Eastern tree frog (*Hyla orientalis*) as our study species. *Hyla orientalis* is a cryptic species of the European tree frog (*Hyla arborea*) group, distributed from the Caspian Sea to the Baltic Sea (Stök et al. 2012). Females start to breed at 2-3 years of age (Özdemir et al. 2012), which means that 10-15 generations have pass since the Chernobyl accident (1986). The species requires warm temperature for the start of the breeding season, which normally occurs in May-June in the study area. *H. orientalis* hibernates buried in the soil or under rocks, leaf litter or wood. Adults feed on a large diversity of small arthropods.

During three consecutive years (2016-2018), we collected adult males of *H. orientalis* actively calling during the breeding season in ponds located within the Chernobyl Exclusion Zone (Ukraine, Figure 1). In total, we examined 226 *H. orientalis* males from twelve localities within the Chernobyl Exclusion Zone (Table 1; Figure 1). Frogs were captured during the night (from 10pm to 1am), placed in plastic bags and transported to our field laboratory in Chernobyl. On the next morning, we recorded different morphological traits of each frog (snout-to-vent length, body depth and width) using a calliper to the nearest 1 mm, and we weighted each individual using a precision balance to the nearest 0.01 g. Morphometric measurements were used to define individual shapes in order to estimate individual dose rates (see below). Once morphometric measurements were recorded, we euthanized frogs by pithing without decapitation (AVMA, 2020), and tissue and bone samples were stored for radiological evaluation. All animals were collected, and procedures conducted, under permit of Ministry of Ecology and Natural Resources of Ukraine (No. 517, 21.04.2016).

### Field estimation of radiation levels

At each locality, we estimated ambient dose rate using a radiometer MKS-AT6130 to measure both gamma dose rate (μSv/h) and the flux of beta particles (counts cm^-2^ min^-1^) at ca. 5 cm above the surface of water (0.3-1.0 m depth) in five random points along the shoreline and in surrounding terrestrial environment. In most cases, the shoreline values had lower variability, while the terrestrial and air (i.e. above water level) values varied substantially. We assume that shoreline values are more indicative of the environment used by frogs during the breeding season, and therefore we used those values for dose assessment (Table 1).

### External exposure: deposits of ^90^Sr and ^137^Cs in the soil of the study localities

In order to estimate radioactive levels of the study localities, and its contribution to external dose rates, we used a spatial database derived from the integration of the airborne gamma survey and the results of soil sampling in earlier 1990s (Arkhipov et al., 1995). The final database represents a geo-positioned 100 × 100 m grid with values of total ^90^Sr and ^137^Cs deposits fell out after the Chernobyl accident. To estimate ^90^Sr and ^137^Cs activity for the sampling localities, we estimated the geometric mean (n= 50 points) from these integrated databases over a 400 meters radius area centred on the study pond, and activity estimates were decay-corrected to the time of the current study (spring 2016-2018; Figure 1). A similar approach, and spatial database, has been previously used in studies of other animals with relatively large home ranges, when direct evaluation of soil sampling was unfeasible (amphibians, Gashchak et al., 2009a; birds, Gaschak et al., 2009b; rodents, Maklyuk et al., 2007; bats Gashchak et al., 2010).

### Internal exposure: estimation of ^90^Sr activity concentration in bones

Relatively high ^90^Sr activity concentration is found in the bones of animals living in the Chernobyl Exclusion Zone (e.g. Maklyuk et al., 2007; Gaschack et al., 2009b, 2010), which allows the application of standard beta spectrometry methods (Bondarkov et al., 2002). In our study, we sampled a femur bone of every frog that was thoroughly cleaned up from remains of soft tissues. Then, we dried the bone sample in order to estimate dry mass to the nearest 0.01 g. After this, we diluted the sample with concentrated HNO_3_ and H_2_O_2_. We evaporated the obtained solution to generate wet salts, followed by the addition of 1M HNO_3_ to standardize the geometry. We used the final solution for beta-spectrometry, and recalculated the obtained data to the dry mass values of each sample. We used a β-spectrometer EXPRESS-01 with a thin-filmed (0.1 mm) plastic scintillator detector, with the software “Beta+” (developed by the Institute of Nuclear Research at the National Academy of Science of Ukraine). This method allows to measure ^90^Sr content in thick-layered samples with a comparable ^137^Cs content (^137^Cs/^90^Sr ratio not exceeding 30:1; Bondarkov et al., 2002). We processed the obtained experimental spectrum using correlations with the measured spectra from OISN-3 standard mixing sources (Applied Ecology Laboratory of Environmental Safety Centre, Odessa, Ukraine; e.g. ^90^Sr+^90^Y, ^137^Cs and the ^90^Sr + ^90^Y, and ^137^Cs combinations), as well as from background. The minimal detectable activity (MDA) was 0.6 Bq per sample. The small mass of the bone samples and the relatively low contamination of frogs from some localities did not allow to estimate ^90^Sr activity concentration above MDA (Supplementary data).

### Internal exposure: estimation of ^137^Cs activity concentration in muscles

In order to estimate ^137^Cs levels, we sampled muscle tissue from frog legs. We measured the wet mass of the muscle sample to the nearest 0.01 g. Then, we diluted the muscle sample with concentrated HNO_3_ and H_2_O_2_. The obtained solution was evaporated to generate wet salts, followed by the addition of 1M HNO_3_ to standardize the geometry. We used the final solution for gamma-spectrometry, and recalculated the obtained data to the wet mass values of each sample. We measured ^137^Cs activity concentrations on the muscle samples using a Canberra-Packard gamma-spectrometer with a high-purity germanium (HPGe) detector (GC 3019). A OISN-1 standard mixed source (^44^Ti/^137^Cs/^152^Eu; Applied Ecology Laboratory of Environmental Safety Centre, Odessa, Ukraine), including epoxy granules (< 1.0 mm) with 1 g cm^-3^ density, was used for calibration. The minimally detectable activity ranged from 0.1 until 0.3 Bq per sample depending on sample mass and radioactivity of the original sample. The small mass of the muscle samples and the relatively low contamination of frogs from some localities did not allow to estimate ^137^Cs activity concentration above MDA (Supplementary data).

### Estimation of individual total dose rates

To estimate total individual dose rates (TDR, in μGy/h) absorbed by each frog during the breeding season, we first estimated whole-body activity of ^90^Sr and ^137^Cs by integrating radionuclide activity concentrations (see above) with body mass of each individual, and considering the relative mass of bones (10%) and muscles (69%, Barnett et al. 2009). We combined radionuclide activity concentrations in frogs, soil, and water with dose coefficients (in μGy/h per Bq per unit of mass). The use of dose coefficients allows transforming radionuclide activity (Bq/kg, Bq/L) into dose rate (μGy/h), and are specific for each radionuclide/organism/ecological scenario combination. Dose coefficients for *H. orientalis* were calculated for internal and external exposure by taking into consideration a theoretical ecologically scenario for the species during a whole breeding period as follows: 8h/day spent on vegetation at >50 cm above ground, 8h/day on the ground, 7h30/day at the water surface, and 30 min/day at the sediment-water interface (soil depth: 10cm; water depth: 100 cm; grass depth: 10 cm; see Giraudeau et al. 2018, for a similar approach). We calculate doses using EDEN v3 IRSN software (Beaugelin-Seiller et al. 2006). For each tree frog, total individual dose rate was calculated by summing internal and external dose rates.

### Statistical analyses

All statistical analyses were conducted in R software (version 3.6.1, R Development Core Team). We log transformed data of all parameters once we added 0.1 unit to each value of ^90^Sr and ^137^Cs dose rates, and to ambient, internal, external, and total dose rate. Using the whole dataset, we conducted mixed-model regressions (lmer function, package lme4 version 1.1-23) to check for the relationships between ambient and total dose rate. In samples collected within localities with ambient dose rate > 1 μS/h, we conducted a mixed-model regression between internal and external dose rate. All regressions included the factor “locality” as random factor. We also conducted linear models to check for differences between localities in total dose rate, and internal-to-external ratio. For data plotting and visualization, we used the function ggplot included in the package ggplot2 (version 3.3.0).

## Acknowledgments

We thank Sergey Gaschack and Yevgenii Gulyaichenko for his invaluable help in the field and on activity rate estimations, and the administrative personal of the Chornobyl Center for Nuclear Safety, Radioactive Waste and Radioecology (Ukraine) for help with research permits and transportation. Clare Bradshaw helped us during the initial stages of the study, and Karine Beaugelin-Seiller during dose rate calculations. This work was supported by the Swedish Radiation Protection Agency-SSM (SSM2018-2038), the FP7-EURATOM COordination and iMplementation of a pan-European instrumenT for radioecology-COMET project (EU-604974), and by Carl Tryggers Foundation (CT 16:344). Carl Tryggers Foundation scholarship (CT 16:344) and Marie Sklodowska-Curie fellowship (METAGE-797879) supported PB, an IRSN doctoral fellowship supported CC, the Institute for Radioecological Protection and Nuclear Safety (IRSN) supported JMB, and the Spanish Ministry of Science, Innovation and Universities (Ramón y Cajal program, RYC-2016-20656) supported GO.

## Author’s Contributions

GO conceived and designed the study; PB, JMB, and GO carried out the field work; CC and JMB performed dose rate calculations; PB analysed the data; PB and GO wrote the paper with inputs from CC and JMB.

## References

Arkhipov, N.P., Voitsekhovich, O.V., Gladkov, G.N., Dzhepo, S.P., Drapeko, G.F., et al., Bulletin of ecological state of the exclusion zone for the first half-year 1995. Ministry of Ukraine for Protection of Public from Consequences of the Accident on Chernobyl NPP. Chernobyl, Issue 5 (in Russian). (1995).

AVMA. Guidelines for the euthanasia of animals: 2020 Edition. American Veterinary Medical Association. Schaumburg, IL. (2020).

Barnett, C. L., Belli, M., Beresford, N. A., Bossew, P., Boyer, P. et al. Quantification of radionuclide transfer in terrestrial and freshwater environments for radiological assessments. IAEA-TECDOC-1616 (2009).

Beaugelin-Seiller, K., Jasserand, F., Garnier-Laplace, J. & Gariel, J. C. Modelling radiological dose in non-human species: principles, computerization, and application. Health Phys. 90, 485–493 (2006).

Beresford, N. A. & Copplestone, D. Effects of ionizing radiation on wildlife: What knowledge have we gained between the Chernobyl and Fukushima accidents? Integr. Environ. Assess. Manag. 7, 371–373 (2011).

Beresford, N. A., Gaschak, S., Barnett, C. L., Howard, B. J., Chizhevsky, I. et al. Estimating the exposure of small mammals at three sites within the Chernobyl exclusion zone – a test application of the ERICA tool. J. Environ. Radioact. 99, 1496–1502 (2008).

Beresford, N. A., Barnett, C. L., Brown, J. E., Cheng, J.-J., Copplestone, D., et al. Predicting the radiation exposure of terrestrial wildlife in the Chernobyl exclusion zone: an international comparison of approaches. J. Radiol. Prot. 30, 341–373 (2010).

Beresford, N. A., Adam-Guillermin, C., Bonzom, J.-M., Garnier-Laplace, J., Hinton, T. et al. Comment on “Abundance of birds in Fukushima as judged from Chernobyl” by Møller et al. (2012). Environ. Pollut. 169, 136 (2012).

Beresford, N., Horemans, N., Copplestone, D., Raines, K. E., Orizaola, G. et al. Towards solving a scientific controversy – The effects of ionising radiation on the environment. J. Environ. Radioact. 211, 106033 (2020a).

Beresford, N. A., Scott, E. M. & Copplestone, D. Field effects studies in the Chernobyl Exclusion Zone: Lessons to be learnt. J. Environ. Radioact. 211, 105893 (2020b).

Beresford, N. A., Barnett, C. L., Gashchak, S., Maksimenko, A., Guliaichenko, E., Wood, M. D. & Izquierdo, M. Radionuclide transfer to wildlife at a ‘Reference Site’ in the Chernobyl Exclusion Zone and resultant radiation exposures. J. Environ. Radioact. 211, 105661 (2020c).

Bondarkov, M. D., Maksimenko, A. M. & Zheltonozhsky, V.A. Non radiochemical technique for 90Sr measurement. Radioprotection 37, C1-927–C1-931 (2002).

Brown, J. E., Alfonso, B., Avila, R., Beresford, N. A., Copplestone, D. et al. The ERICA tool. J. Environ. Radioact. 99, 1371–1383 (2008).

Burraco, P., Car, C., Bonzom, J.-M., Beaugelin-Seiller, K., Gaschack, S. & Orizaola, G. Lack of impact of exposure to radiation on blood physiology biomarkers of Chernobyl tree frogs. Front. Zool. 18, in press (2021).

Garnier-Laplace, J., Copplestone, D., Gilbin, R., Alonzo, F., Ciffroy, P. et al. Issues and practices in the use of effects data from FREDERICA in the ERICA Integrated Approach. J. Environ. Radioact. 99, 1474–1483 (2008).

Gashchak, S. P., Beresford, N. A., Maksimenko, A. M. & Vlaschenko, A. S. Strontium-90 and caesium-137 activity concentrations in bats in the Chernobyl exclusion zone. Radiat. Environ. Bioph. 49, 635–644 (2010).

Gashchak, S. P., Maklyuk, Y. A., Maksimenko, A. M. & Bondarkov, M. D. Radioecology of amphibians in Chernobyl zone. Problems of the Chernobyl Exclusion Zone 9, 76–86 (in Russian) (2009a).

Gaschak, S., Bondarkov, M., Makluk, J.U., Maksimenko, A., Martynenko, V., et al. Assessment of radionuclide export from Chernobyl zone via birds 18 years following the accident. Radioprotection 44, 849–852 (2009b).

Geras’kin, S. A., Fesenko, S.V. & Alexakhin, R. M. Effects of nonhuman species irradiation after the Chernobyl NPP accident. Environ. Int. 34, 880–897 (2008).

Giraudeau, M., Bonzom, J.-M., Ducatez, S., Beaugelin-Seiller, K., Deviche, P., et al. Carotenoid distribution in wild Japanese tree frogs (*Hyla japonica*) exposed to ionizing radiation in Fukushima. Sci. Rep. 8, 7438. (2018).

ICRP. Environmental protection: the concept and use of reference animals and plants. Annals of the ICRP 38, 1–242 (2008).

Intelligence Systems GEO. Atlas of radioactive contamination of Ukraine. Ministry of Emergencies and Affairs of Population Protection from the Consequences of Chernobyl Catastrophe, Ukraine (2011).

Leadon S. A. Repair of DNA damage produced by ionizing radiation: a minireview. Semin. Radiat. Oncol. 6, 295–305 (1996).

Maklyuk, Y. A., Maksimenko, A. M., Gaschak, S. P., Bondarkov, M. D. & Chizhevsky, I. V. Long-term dynamic of radioactive contamination (90Sr, 137Cs) of small mammals in Chernobyl zone. Ecology 38, 198–206 (in Russian) (2007).

Møller, A. P. & Mousseau, T. A. Biological consequences of Chernobyl: 20 years on. Trends Ecol. Evol. 21, 200–207 (2006).

Møller, A. P. & Mousseau, T. A. Are organisms adapting to ionizing radiation at Chernobyl? Trends Ecol Evol. 31, 281–289 (2016).

Özdemir, N., Altunişik, A., Ergül, T., Gül, S., Tosunoğlu, M. et al. Variation in body size and age structure among three Turkish populations of the treefrog *Hyla arborea*. Amphibia-Reptilia 33, 25–35 (2012).

Raines, K. E., Whitehorn, P. R., Copplestone, D. & Tinsley, M. C. Chernobyl-level radiation exposure damages bumblebee reproduction: a laboratory experiment. Proc. R. Soc. B. 287, 20201638 (2020).

Santivasi, W. L. & Xia, F. Ionizing radiation-induced DNA damage, response, and repair. Antioxid. Redox Signal. 21, 251–259 (2014).

Sahoo, S. K., Kavasi, N., Sorimachi, A., Arae, H., Tokonami, S. et al. Strontium-90 activity concentration in soil samples from the exclusion zone of the Fukushima daiichi nuclear power plant. Sci. Rep. 6, 23925 (2016).

Sohrabi M. World high background natural radiation areas: Need to protect public from radiation exposure. Radiat. Meas. 50, 166–171 (2013).

Stark, K., Avila, R. & Wallberg, P. Estimation of radiation doses from 137Cs to frogs in a wetland ecosystem. J. Environ. Radioact. 75, 1–14 (2004).

Stöck, M., Dufresnes, C., Litvinchuk, S. N., Lymberakis, P., Biollay, S. et al. Cryptic diversity among Western Palearctic tree frogs: Postglacial range expansion, range limits, and secondary contacts of three European tree frog lineages (*Hyla arborea* group). Mol. Phylogenet. Evol. 65, 1–9 (2012).

