## Supplementary figures and images for "Assessment of exposure to ionizing radiation in Chernobyl tree frogs (*Hyla orientalis*)"

### Figure S1

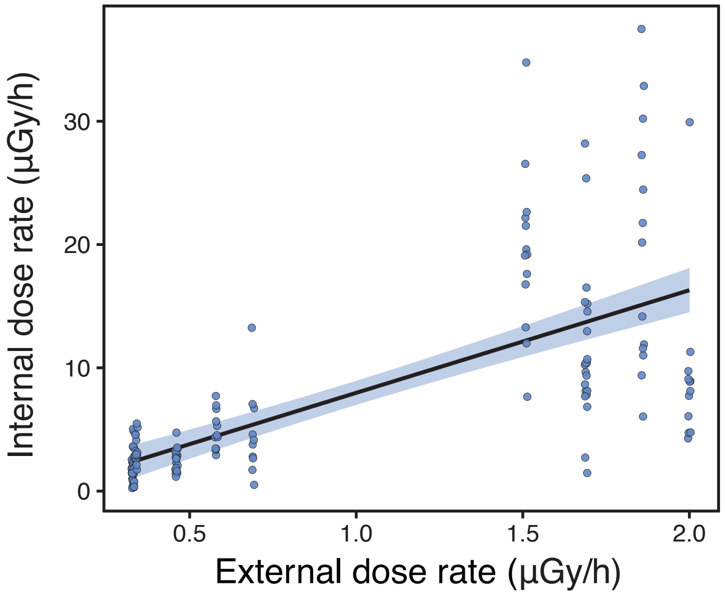

### Figure S2

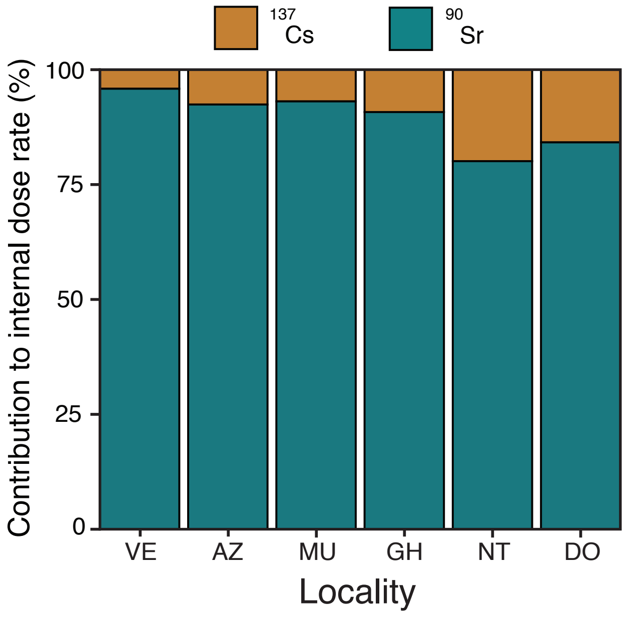

### Figure S3

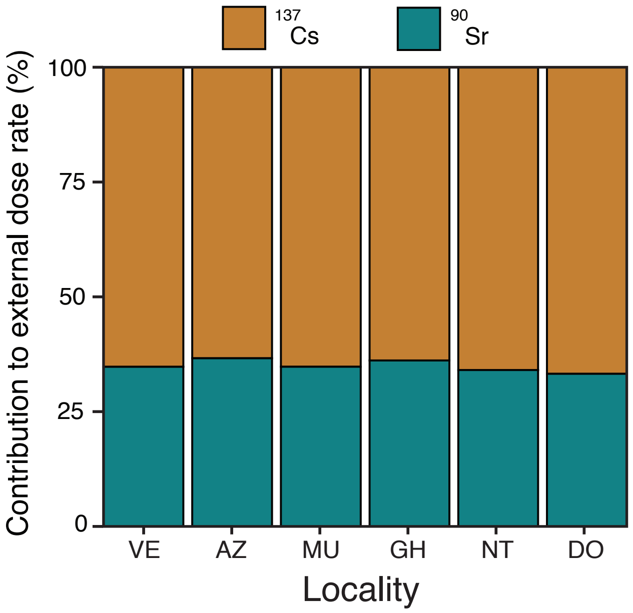
